# Genetic structure in patchy populations of a candidate foundation plant: a case study of *Leymus chinensis* (Poaceae) using genetic and clonal diversity

**DOI:** 10.1101/2021.06.12.448174

**Authors:** Jian Guo, Christina L. Richards, Kent E. Holsinger, Gordon A. Fox, Zhuo Zhang, Chan Zhou

## Abstract

**PREMISE:** The distribution of genetic diversity on the landscape has critical ecological and evolutionary implications. This may be especially the case on a local scale for foundation plant species since they create and define ecological communities, contributing disproportionately to ecosystem function.

**METHODS:** We examined the distribution of genetic diversity and clones, which we defined first as unique multi-locus genotypes (MLG), and then by grouping similar MLGs into multi-locus lineages (MLL). We used 186 markers from inter-simple sequence repeats (ISSR) across 358 ramets from 13 patches of the foundation grass *Leymus chinensis*. We examined the relationship between genetic and clonal diversities, their variation with patch-size, and the effect of the number of markers used to evaluate genetic diversity and structure in this species.

**RESULTS:** Every ramet had a unique MLG. Almost all patches consisted of individuals belonging to a single MLL. We confirmed this with a clustering algorithm to group related genotypes. The predominance of a single lineage within each patch could be the result of the accumulation of somatic mutations, limited dispersal, some sexual reproduction with partners mainly restricted to the same patch, or a combination of all three.

**CONCLUSIONS:** We found strong genetic structure among patches of *L. chinensis*. Consistent with previous work on the species, the clustering of similar genotypes within patches suggests that clonal reproduction combined with somatic mutation, limited dispersal, and some degree of sexual reproduction among neighbors causes individuals within a patch to be more closely related than among patches.

## INTRODUCTION

Genetic diversity provides the raw material for evolutionary change and thus the potential to adapt to changing environments (Frankel and Soulé, 1981), but how genetic diversity is distributed on the landscape has critical ecological and evolutionary implications (Slatkin, 1987; Whitham et al., 2006; Hughes et al., 2008; Sork et al., 2013; Hughes and Lotterhos, 2014). Although the theoretical importance of genetic diversity for the long-term persistence of populations is well understood, we have little information about the fine-scale distribution of genetic variation in natural populations (Morris et al., 2011; Gray et al., 2014; Hughes and Lotterhos, 2014). This is true even for species that are quite common or dominate a landscape. According to Ellison (2019), 50 years of research has identified that there are “foundation species” that 1) dominate a community of species (numerically and in overall size), 2) determine the diversity of associated taxa through non-trophic interactions, and 3) modulate fluxes of nutrients and energy in their ecosystem. Although it is challenging to document that a candidate species meets all of the requirements to be considered a foundation species, there is growing recognition that protection and restoration of these species can disproportionately maintain habitat integrity and ecosystem resilience (Keith et al., 2017; Ellison, 2019; Qiao et al., 2021). The genetic make-up of foundation plant species can be particularly important on a local scale since these species create and define ecological communities and contribute disproportionately to ecosystem function (Ellison et al., 2005; Hughes and Lotterhos, 2014; Lau et al., 2016; Keith et al., 2017; Whitham et al., 2020).

Grassland species are often able to spread clonally through apomictic or vegetative means. In spite of their ecological importance in these systems, clonal organisms are thought to be limited in evolutionary potential since they lack the ability to generate variation and purge deleterious mutations through recombination (Lynch et al., 1993; Holsinger, 2000; but see Verhoeven and Preite, 2014). However, most clonal plants have some level of sexual reproduction (Richards, 1986), and an estimated two-thirds of temperate plant species are thought to be capable of clonal propagation (Van Groenendael and De Kroon, 1990; Klimes et al., 1997; Schoen and Schultz, 2019). Genetic diversity at marker loci is commonly measured as the proportion of polymorphic loci and as expected heterozygosity among all individuals, while clonal (or genotypic) diversity can be defined by the number of unique genotypes (evaluated across multiple loci) within a group (Sole et al., 2004; Arnaud-Haond et al., 2007; Reynolds et al., 2012). Although functional genetic variation is most important for understanding evolutionary responses (Texeira and Huber, 2021), the distribution of ramets per clone provides information on how genetic diversity is structured and therefore how natural selection might be constrained (Frankel, 1970; Widén et al., 1994; Arnaud-Haond et al., 2007). Hence, when considering how genetic diversity is spatially structured in plants that can spread vegetatively, it is useful to consider the abundance and distribution of ramets per clone in addition to population levels of genetic diversity. When conserving or restoring communities of such clonal plants, the relationship between genetic diversity and clonal diversity is an important consideration. The two metrics capture different aspects of genetic structure that may be related to the adaptive capacity and long-term viability of these plants (Schaal et al., 1991; Holsinger, 2000; Han et al., 2007; Reynolds et al., 2012). Several studies have found that populations of clonal species are generally just as diverse as non-clonal species (Ellstrand and Roose, 1987; Widén et al., 1994; Franks et al., 2004; Houliston and Chapman, 2004; Keeler et al., 2002; Richards et al., 2004; Raabová et al., 2015), but there are important exceptions particularly in the context of range expansion or invasive species (Ainouche et al., 2004, 2009; Gao et al., 2010; Richards et al., 2012; Zhang et al., 2016; Mounger et al., 2021).

Clonal foundation plant species can form vast monospecific stands, but they may also be found in patches that vary greatly in size in the wild due to disturbance, patch age and the favorability of conditions (Daehler and Strong, 1994; Davis et al., 2004; Travis and Hester, 2005; Morris et al., 2011; Gray et al., 2014; Hughes and Lotterhos, 2014). Genetic structure in clonal plants partly results from the fact that a single individual that establishes by seed or vegetative organ can produce many individual ramets through clonal propagation (Richards, 1986; Lehmann, 1997; Chen et al., 2006), which can exhibit a patchy, interdigitated distribution (Zhou et al., 2003; Franks et al., 2004; Richards et al., 2004). Ramets that belong to a given genet can also accumulate mutations and appear to be new MLGs (Schoen and Schulz, 2019), even though functionally they could be more appropriately considered the same lineage (hence MLL). This can lead to different patterns of genetic diversity in older patches. In one scenario, patches may accumulate a large number of lineages, which each expand to result in a large patch area with high clonal diversity (Raabová et al., 2015). On the other hand, patches may have limited recruitment, or competitively superior lineages may produce many ramets through vegetative propagation to enlarge the patch area such that long-lived populations may be characterized by fewer genotypes (Sakai, 1995; Gardner and Mangel, 1999; Travis and Hester, 2005). However, detecting these different patterns of clonal structure can be complicated by outcrossing between closely related lineages, as well as the fact that somatic mutations may accumulate and provide an important source of variation in older lineages (Schoen and Schultz, 2019).

Despite the fact that most of evolutionary theory and models are based on sexually reproducing, genetically distinct individuals, with the rapidly escalating application of molecular approaches to ecological questions, researchers have developed some sophisticated approaches to more accurately estimate genetic and clonal diversity in organisms that have a mix of both types of reproduction (Arnaud-Haond et al., 2007; Kamvar et al., 2014, 2015). In particular, it is useful to define a multi-locus lineage (MLL) as a set of distinct multi-locus genotypes (MLGs) that are more closely related to one another than any of them are to other genotypes. Identifying unique MLLs requires enough polymorphic markers to provide discriminating power to differentiate closely related genotypes, but also requires identification of scoring errors that will make estimates of relationships inaccurate (Douhovnikoff and Dodd, 2003; reviewed in Arnaud-Haond et al., 2007). In the past, researchers did not distinguish between genetically distinct individuals (MLGs) and MLLs (e.g., Franks et al., 2004; Richards et al., 2004). As the number of markers has increased, however, allowing more closely related MLGs to be distinguished from one another, it has also become more important to distinguish among degrees of relationship (Douhovnikoff and Dodd, 2003; Arnaud-Haond et al., 2007; Kamvar et al., 2015). A common approach to identifying degrees of relationship is to use clustering algorithms to identify MLLs, groups of genotypes that are more similar to one another than any of them is to another genotype (Kamvar et al., 2015).

In this study, we examined the relationship between patch size and genetic structure in patches of the clonal plant *Leymus chinensis,* a dominant perennial grass species in the Chinese steppe zone (Li et al., 2014; Zhou et al., 2014; Zhou et al., 2015). While this species has not been formally referred to as a foundation species, several authors have argued that *L. chinensus* is one of the most important dominant species of the undisturbed mature community native to the Inner Mongolia grasslands (Bai et al., 2004, 2010; Tong et al., 2004; Meng et al., 2019). Previous work comparing undisturbed mature grassland communities to severely degraded communities showed that *L. chinensis* had the highest relative biomass and lowest interannual variability of biomass production among all species in the undisturbed system which contributed to ecosystem stability (Bai et al., 2004). These studies suggest that *L. chinensis* is a good “candidate foundation species” which will require further validation (sensu (Qiao et al., 2021)). Only a few studies have examined genetic variation in *L. chinensis*, and they have focused on large geographical scales, finding significant variation among different regions of China (Wang et al., 2005) and significantly higher genetic diversity in eastern than in western sites along a longitudinal gradient in northeast China (Yuan et al., 2016). However, there is very little information about how the genetic diversity of *L. chinensis* is distributed among different patches on a micro-geographical scale (e.g., Hong, 2013) or how genetic diversity is related to ecosystem resilience in this system (e.g., Bau et al., 2004). This knowledge is essential for managers to formulate the best strategy for conservation and restoration of this important grassland species. The structure of genetic diversity in patchy populations of *L. chinensis* could be an unappreciated but important component of successful restoration of degraded lands that were once dominated by this species (Bai et al., 2004; Tong et al., 2004). Using a large number of Inter Simple Sequence Repeat (ISSR) markers, we addressed the following questions: (1) What is the relationship between genetic and clonal diversity? (2) Do larger patches have higher levels of genetic and clonal diversity? (3) Is genetic or clonal structure correlated with patch size? Because we used a large number of ISSR markers, we also explored the technical question of how the number of markers used affects conclusions about the numbers of MLLs, as well as whether patches are predominantly composed of MLGs more closely related to one another than to MLGs in other patches.

## MATERIALS AND METHODS

### Study species

*Leymus chinensis* is a dominant tetraploid perennial grass species in the Chinese steppe zone (Li et al., 2014; Zhou et al., 2014; Zhou et al., 2015) occurring on a total area in Asia of approximately 420,000 km^2^, of which 220,000 km^2^ are located in China (Inner-Mongolia and Ning Xia Investigation and Survey Team of the Chinese Academy of Sciences 1985). *Leymus chinensis* is a significant forage grass, with high nutrient value, good palatability and high above-ground biomass production (Zhu, 2004; Bai et al., 2009). However, vast acreage of the species has suffered from overgrazing which has led to “steppe degradation” (Bai et al., 2004; Tong et al., 2004). As an ecologically and economically important foundation grass, *L. chinensis* has received considerable attention in recent years (Bai et al., 2004; Tong et al., 2004; Zhu, 2004).

A large number of investigations have focused on growth (Li et al., 2014; Liu et al., 2011), reproduction (Bai et al., 2009) and physiology (Xu and Zhou, 2007) of the species. *Leymus chinensis* can successfully survive under stressful conditions, such as drought, salinity, alkalinity, and depleted organic matter. The species possesses the capacity for both vegetative propagation through rhizome fragmentation and sexual reproduction by seeds (Bai et al., 2009). In natural habitats, *L. chinensis* appear in differently sized patches, and a single genet can give rise to multiple ramets and extensively spreading clones (Yang et al., 2006). Vegetative reproduction is thought to be more common than sexual reproduction (Wang, 2001; Zhang et al., 2005), but the actual contributions of these two modes of reproduction is unknown. When it reproduces sexually, *L. chinensis* is partially self-incompatible (with self-pollen leading to seed set in 5-10% of experimental pollinations, as compared with 60-84% of non-self-pollen) (Zhang et al., 2004); and as in other grasses the SI system is gametophytic (Chen et al., 2019).

### Study sites

This study was carried out at the Grassland Ecological Research Station of Northeast Normal University, Changling County, Jilin Province, China (44°30′ to 44°45′N, 123°31′ to 123°56′E). This area is in the northeast of the Chinese steppe zone. The grassland has low-lying topography, and a typical mesothermal monsoon climate with hot, rainy summers and cold, arid winters. Mean annual precipitation ranges from 300 to 450 mm, more than 60% of which occurs from June to September. Mean annual temperature ranges from 4.6 to 6.4°C and the frost-free period lasts for 130 to 165 days (Guo et al., 2020). The soil is a mixed salt-alkali meadow steppe (Salid Aridisol, US Soil Taxonomy) with 29% sand, 40% silt and 31% clay (top 10 cm) (Zhu, 2004). Grazing of this area was prohibited beginning in September 1995. Other plant species typically co-occuring with *L. chinensis* include *Phragmites communis*, *Kalimeris integrifolia*, *Carex duriuscula*, *Calamagrostis epigeios*, and *Chloris virgata*.

### Sampling design

We collected tissue samples in August 2010, when the plants were vigorously growing. Thirteen *L. chinensis* patches were selected haphazardly within an area of 2000m^2^. Patches were defined naturally (by the space occupied by *L. chinensis* locally), and all patches were separated from others by at least 200m. Patches ranged in area between 2.8m^2^ and 166.8m^2^ We collected vegetative tillers of *L. chinensis* at distances of 0, 0.5, 1, 2, 4, 8, and 16m from the center of each patch (or to the edge of the patch, whichever came first) in each of eight directions (N, S, E, W, NE, NW, SE, SW). Consequently, we sampled between 17-35 ramets from each patch (Table 2). The vegetative ramets of *L. chinensis* were harvested at the ground surface and each ramet was put into a zip-lock plastic bag with silica gel to preserve the tissue. We transported all samples to the laboratory and stored them at −20°C.

### DNA extraction

For each sample, we performed extractions of total genomic DNA using a modified CTAB method (Doyle and Doyle, 1987). We ground tissue of dried young leaves (0.025 to 0.030g) to a fine powder in liquid nitrogen with pure quartz sand in a 2 ml centrifuge tube. Then, we added 400 μl of extraction buffer [2% cetyl trimethyl ammonium bromide (CTAB); 100 mM Tris (pH=8.0); 20 mM ethylenediamine tetra-acetic acid (EDTA); 0.5% NaHSO_3_; 1.4 M NaCl; 1% polyvinylpyrrolidone (PVP); 2% β-mercaptoethanol] to the ground sample and incubated at 65°C for 30 minutes. We performed two rounds of extraction with 400 μl of tris-saturated phenol. After centrifugation, we performed two more rounds of extraction with chloroform: isoamyl alcohol (24:1) and precipitated the DNA with 200 μl of 5 M potassium acetate and 1ml of 100% ice-cold ethanol. The precipitate was washed with 70% ethanol twice, dried, and resuspended in 100ul of 1×Tris-EDTA (TE) buffer (Shimizu et al., 2002). We estimated the quantity of DNA by comparing band intensities with known amounts of lambda DNA on 0.8% agarose gel. DNA quality was checked by spectrophotometry at 260 nm and 280 nm. All the DNA samples were stored at −20°C for ISSR analysis.

### ISSR amplification

Twenty-one primers from the Biotechnology Laboratory, University of British Columbia (UBC set no.9) were synthesized by Sangon (Shanghai Sangon Biological Engineering Technology and Services Co. Ltd, Shanghai, China) for initial screening. Eventually, we selected 11 primers that produced clear bands with good reproducibility and high polymorphism (Table 1).

**TABLE 1.** Basic properties of the 11 ISSR primers producing 186 polymorphic loci: annealing temperature in PCR (T_A_), number of bands scored (N-bands): 100% of scored bands were polymorphic.

| Primer | Primer sequence (5'-3') | $T_A$ (°C) | N-bands |
| --- | --- | --- | --- |
| **6** | HVH(CA)<sub>7</sub>T | 58 | 19 |
| **10** | (CA)<sub>8</sub>RG | 58 | 14 |
| **13** | (AC)<sub>8</sub>YG | 56 | 16 |
| **807** | (AG)<sub>8</sub>T | 54 | 13 |
| **811** | (GA)<sub>8</sub>C | 56 | 23 |
| **818** | (CA)<sub>8</sub>G | 58 | 15 |
| **827** | (AC)<sub>8</sub>G | 56 | 18 |
| **834** | (AG)<sub>8</sub>YT | 54 | 14 |
| **842** | (GA)<sub>8</sub>YG | 55 | 15 |
| **848** | (CA)<sub>8</sub>RG | 58 | 19 |
| **855** | (AC)<sub>8</sub>YT | 57 | 20 |

We performed ISSR amplification in a volume of 10 μl containing 10ng template DNA, 1μl 10×PCR Buffer, 20 mM MgCl_2_, 10 mM dNTPs, 1.5 μM primer, and 0.4U of Taq DNA polymerase. All amplifications were carried out in a Veriti 96 Well Thermal Cycler (Applied Biosystems, USA). For primers 6, 10, 13, 807, 818, 827, 834, 842, 848 and 855, the PCR reaction procedure was as followed: an initial denaturation step of 2 min at 94°C, 35 cycles of 1 min at 94°C, 1 min at respective *T*_m_ values (Table 1), and 1 min at 72°C, and a final extension step of 10 min at 72°C. For primer 811, the amplifications began with initial denaturation for 5 min at 94°C, followed by 45 cycles of 30 s at 94°C, 45 s at 56°C, and 2 min at 72°C, and a final extension at 72°C for 5 min. The amplification products obtained were separated on 2% agarose gels containing 0.1% ethidium bromide at 120 V for 2h, and photographed under UV light. The molecular weight of every amplified band was evaluated by using the DL 2000 Marker (Beijing Dingguo Changsheng Biotechnology Co. Ltd, Beijing, China). Each reaction was repeated twice for each sample with each primer. Only those primers which generated reproducible bands in both reactions for each sample were used for the data analysis.

### Data analysis

Only fragments that were distinct, reproducible, and well-resolved were included in the analysis. We scored all amplified fragments as present (1) or absent (0) by comparing size with external standards.

We used R software, version 3.43 (R Core Team, 2017), for all data analyses unless otherwise noted. We examined graphically the richness and distribution of markers within each of the 13 patches (Figure 1, Table 2), and generated a heatmap of marker frequencies using the heatmap.2 function in the R library gplots. For purposes of our analysis we define a multilocus lineage as a set of distinct genotypes more closely related to one another than any of them are to other genotypes. To assess the number of multilocus lineages (MLLs) we used three approaches: (1) we examined the density plot of numbers of marker differences between all pairs of sampled individuals, to seek an intermediate minimum (sensu Douhovnikoff and Dodd, 2003; Kamvar et al., 2014, 2015). If the distribution of markers is the consequence of differentiation between MLLs (to the right) and somatic mutation within lineages plus measurement error (to the left), the minimum provides a criterion for distinguishing genotypes that belong to the same MLL (those with differences to the left of the minimum,) and those belonging to different MLLs (those with difference to the right). Specifically, following Kamvar et al. (2014, 2015) we calculated the Prevosti distances between all pairs of samples and used these distances to estimate a threshold to distinguish within-lineage from among-lineage differences. (2) we assigned ramets to MLLs using the infinite alleles model in GenoDive (Meirmans and Van Tienderen, 2004; Meirmans, 2020), using cutoffs between 14 and 42 marker differences (based on the approximate location of a minimum in the density plot) to determine clonality. We then examined the resulting estimates of clonal diversity in each patch as the threshold varied, as an independent estimate of the threshold for distinguish within- from among-lineage diversity. (3) We used an assignment test (Meirmans, 2020) to determine whether ramets were appropriately assigned to patches of origin.

**FIGURE 1.**
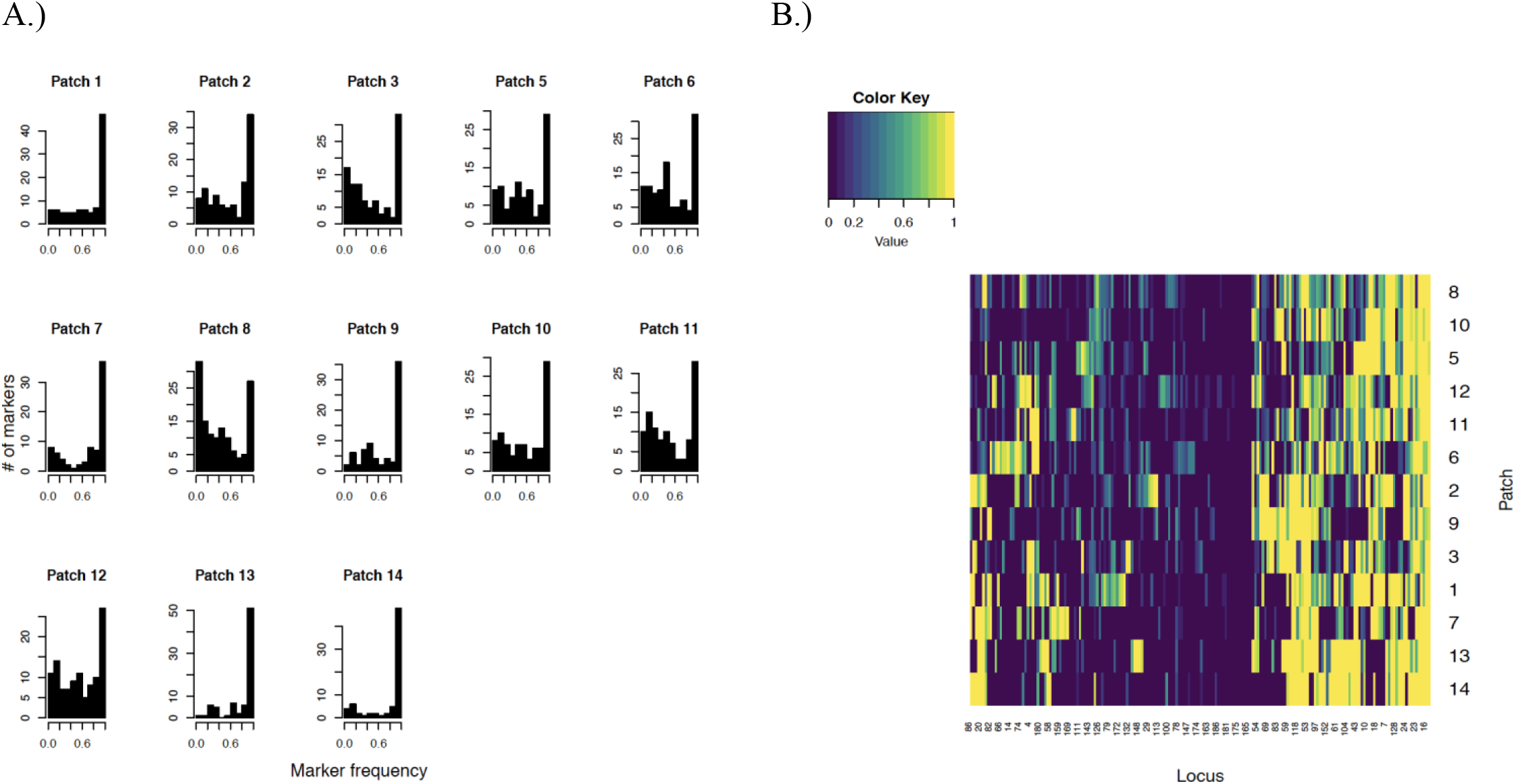
A). Total Number ISSR markers that occur in a given frequency, by patch. B.) Relative frequencies of individuals in each patch carrying at least one copy of each locus.

**TABLE 2.** Patch area, numbers of ramets sampled, ISSR markers found (richness), number of polymorphic loci, and percent polymorphic loci (PL) by patch.

| **Patch ID** | **Area (m<sup>2</sup>)** | **N ramets** | **Marker richness** | **N polymorphic loci** | **PL (%)** |
| --- | --- | --- | --- | --- | --- |
| **1** | 13.98 | 26 | 98 | 56 | 57 |
| **2** | 57.70 | 33 | 100 | 62 | 62 |
| **3** | 69.81 | 35 | 103 | 75 | 73 |
| **5** | 8.40 | 26 | 93 | 66 | 71 |
| **6** | 97.23 | 34 | 112 | 85 | 76 |
| **7** | 8.64 | 25 | 78 | 44 | 56 |
| **8** | 166.79 | 35 | 134 | 108 | 81 |
| **9** | 35.57 | 29 | 75 | 45 | 60 |
| **10** | 22.11 | 27 | 87 | 62 | 71 |
| **11** | 13.77 | 26 | 103 | 72 | 70 |
| **12** | 19.29 | 27 | 109 | 84 | 77 |
| **13** | 3.24 | 18 | 80 | 33 | 41 |
| **14** | 2.78 | 17 | 73 | 29 | 40 |
| **Total** |  | 358 | 186 | 186 | 100 |

We report clonal diversity as the proportion of distinguishable genets (the number of genotypes divided by the total number of individuals or G/N). We examined other aspects of diversity using diversity statistics derived from estimates of population structure provided by the R package Hickory v0.1 (https://kholsinger.github.io/Hickory/), which uses Hamiltonian Monte Carlo as implemented in Stan to provide Bayesian estimates of the relevant parameters (including the entire posterior distribution of allele frequencies, within patch inbreeding (*F*_IS_), and diversity within and among patches with the fixation index (*F*_st_)). Hickory implements an improved version of a Bayesian model first described in Holsinger et al. (2002) and Holsinger and Wallace (2004) that addresses shortcomings noted by Foll et al. (Foll et al., 2008). We use custom R scripts to extract the posterior distribution of allele frequencies and use them to provide estimates of several estimates of diversity.

1. We estimate Nei’s Genetic Diversity within each population as 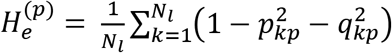 where *p_kp_* and *q_kp_* are the frequency of dominant and recessive alleles at locus *k* in population *p* and *N_l_* is the number of loci. We do not apply the bias correction typically used in estimates of *H_e_*, because the posterior estimates of allele frequencies already account for sampling uncertainty.
2. We estimate the number of effective alleles per locus (Kimura and Crow, 1964) as 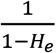.
3. We estimate the Shannon-Weaver allelic diversity as 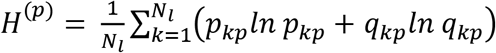.
4. Jost (2006) pointed out that a simple transformation of the Shannon-Weaver diversity provides a second index of the effective number of alleles. We estimate the effective number of alleles derived Shannon-Weaver diversity is exp(*H*^(*p*)^).

In addition to these estimates of genetic diversity, we also estimated the composite disequilibrium ((Weir, 1996); 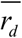) because the marker phenotypes cannot be resolved as alleles. Substantial composite disequilibrium is expected as a result of clonal reproduction.

We examined similarity of individuals and genets within and among patches, using both a clustering approach and neighbor-joining trees. To determine the number of clusters to be used, we used an iterative procedure suggested by Grunwald et al. (2016): we iteratively fit a series of principal component analyses (PCA) specifying the use of between 1 and 15 clusters, based on the Prevosti distances. At each of these steps, we sum the within-cluster sums of squares (SS); we chose the number of clusters at which the sum of SS leveled off at an approximate minimum. Within the R package poppr ver. 2.8.2 (Kamvar et al., 2014, 2015), we used the nj() function to make the neighbor-joining trees and the find.clusters function to identify clusters using the principal components data. Finally, to examine whether diversity measurements were related to patch area or patch sample size, we used Pearson’s correlations.

We used rarefaction methods to evaluate the effect of the number of markers used to identify the number of MLLs in each patch. To do so we resampled our data into multiple unique sets of 10, 30, 50, 70, 90, 110, 130 and 150 markers and then calculated the mean and variance of the number of MLL detected at each number of markers. We then used graphical analysis to examine the relationship between the number of markers used and the means and variances of the estimated numbers of MLLs.

## RESULTS

We found a total of 186 reproducible loci across 358 individuals using 11 ISSR primers. The number of bands amplified by each primer set ranged from 13 to 23, and all amplified loci were polymorphic (Table 1). Within each patch, the number of ISSR bands (marker richness) varied from 73 in patch 14 to 134 in patch 8, with differences largely related to sample size (Pearson’s *ρ* = 0.71) associated with differences in patch area. Similarly, the number of polymorphic loci within patches varied considerably, from 29 in patch 14 to 108 in patch 8 (Table 2).

### Genotypic and genetic diversity

Every one of the 358 individuals sampled possessed a unique multi-locus genotype (MLG): the number of genets per ramet was 1 based strictly on detected polymorphisms. However, patterns of marker abundance varied considerably among patches. Histograms of the frequency of marker occurrence by patch, show that on the order of 25-50 markers occurred in nearly all individuals within a patch (Fig. 1A). Beyond that, patterns of occurrence varied among patches; in several patches (most notably in patch 8) there was a second mode for markers that were very uncommon in that patch. The heatmap of marker presence (Fig. 1B) shows that many markers are quite uncommon (while others are very common) in these samples, and the patches appear to vary in their patterns of relative occurrences of these markers. Overall, we found that many markers occur in over half of the ramets (42 markers, or 23%, occurred in at least half of individuals; Fig. 2A), while a few markers occur in very few ramets (three markers occurred in only one individual and five occurred in only two). A comparison of the number of differences between pairs of ramets indicates that ramets differ by between 1 and 89 markers with 99.4% of comparisons differing by more than 10 markers (Fig. 2B).

**FIGURE 2.**
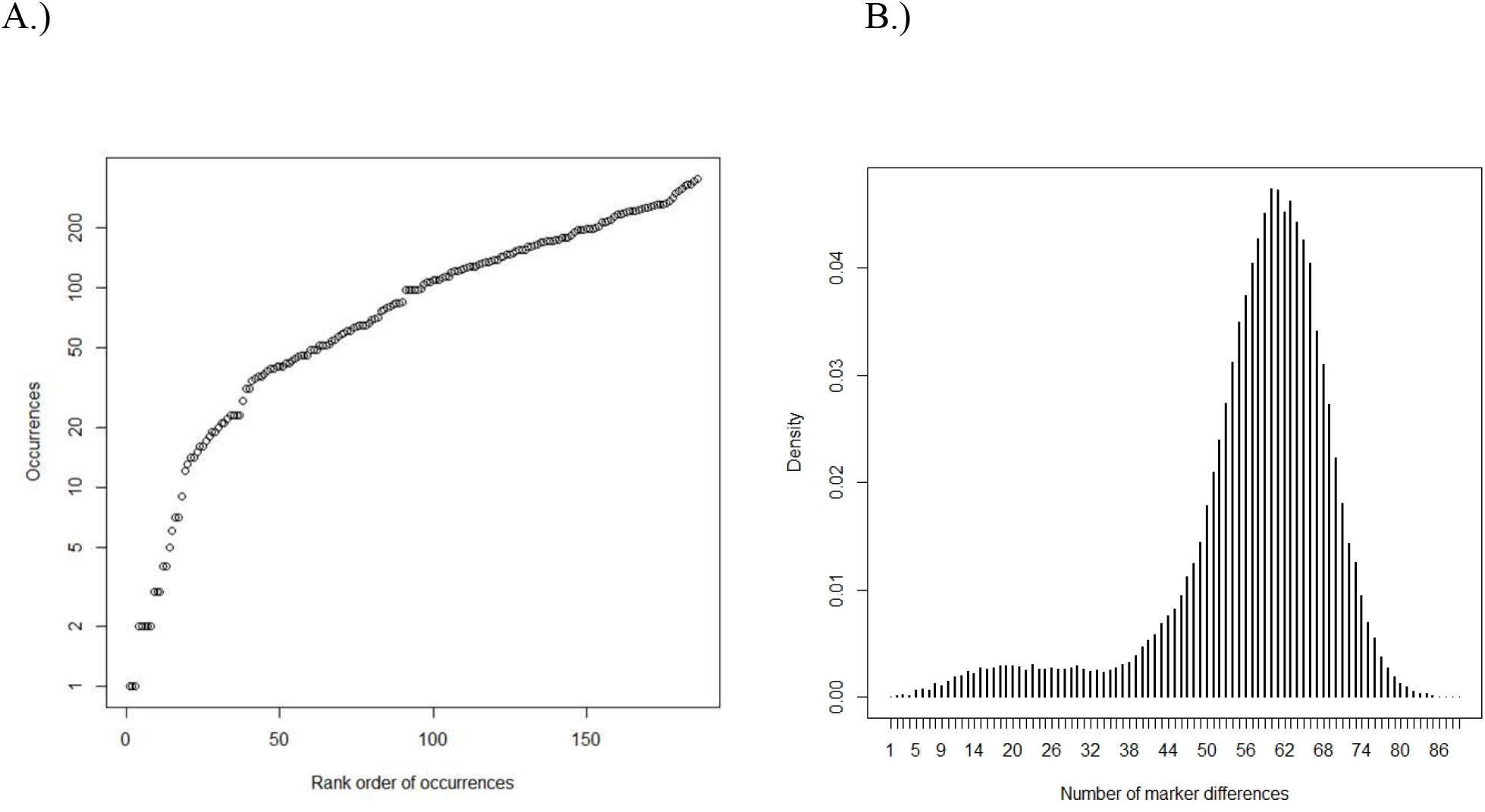
Distribution of 186 ISSR markers across the 358 individuals. A.) Number of occurrences of each of the markers (log scale). B.) Comparisons of number of marker differences between pairs of ramets.

There is a minimum in the density of pairwise differences at approximately 30 differences (Fig. 2B). This corresponds to a Prevosti distance of 0.064 using the farthest neighbor algorithm (Fig. 3). Using this threshold leads to identification of 13 MLLs across the 358 individuals. Other algorithms (nearest neighbor and UPGMA) gave smaller thresholds that had no correspondence with the approximate minimum in the density plot. Henceforth, we refer to the original (N=358) data set as the multi-locus genotype (MLG) data set and the reduced (N=13) data set as the MLL data set. This assignment to MLLs indicated that each patch corresponds to a single genetic lineage, with the exception of patch 8. This patch contains 1 ramet assigned to MLL 2 (otherwise found only in patch 13) and 34 ramets assigned to cluster 13 (found nowhere else).

**FIGURE 3.**
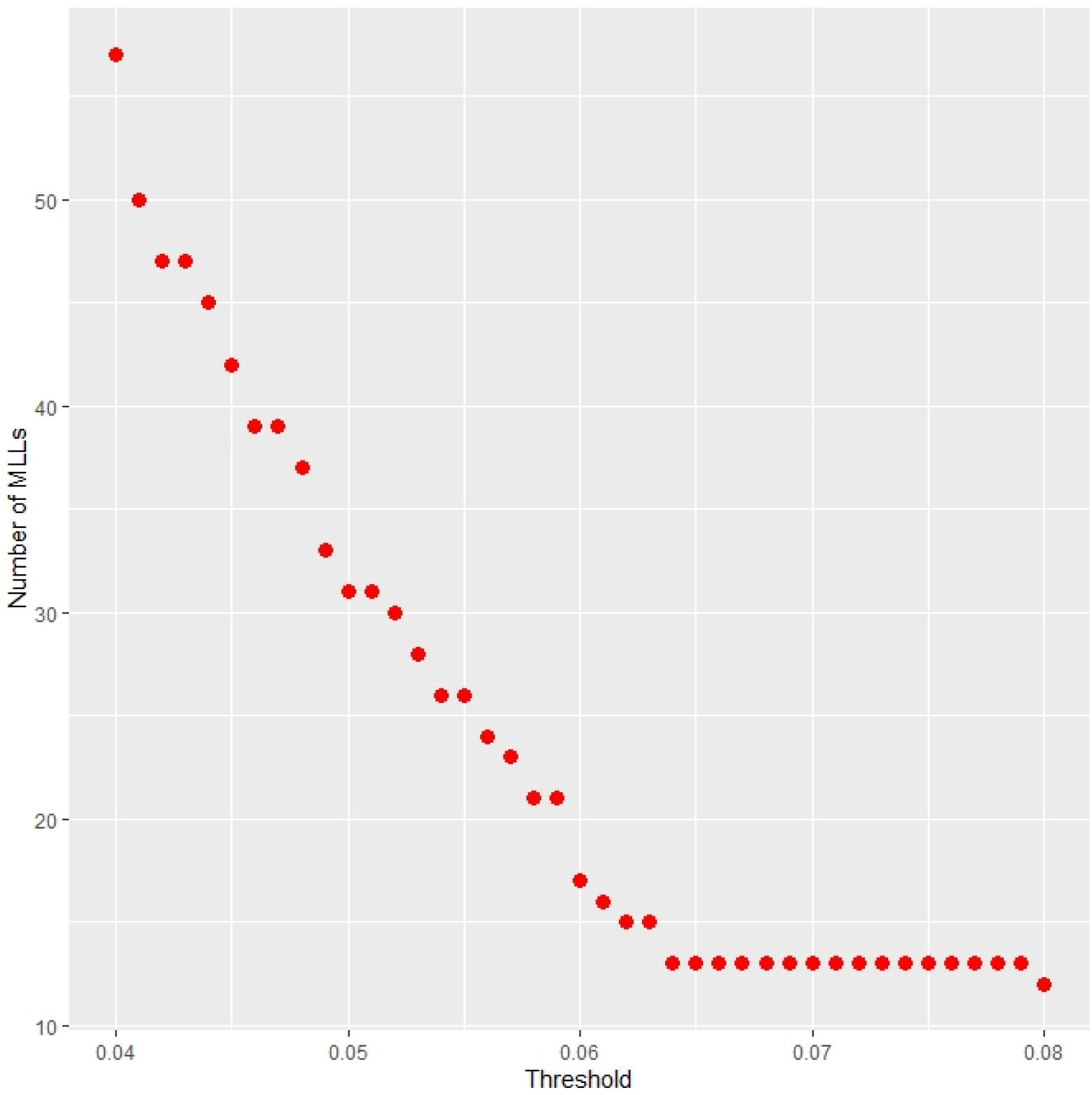
Inferred number of multilocus lineages (MLLs) as a function of the threshold Prevosti distance used to assign ramets to MLLs. For distances less than 0.064, there is a linear decline in inferred number of multilocus lineages (MLLs) with increasing distance. For thresholds between 0.064 and 0.08, there is a stable estimate of 13 MLLs, suggesting 0.064 as an appropriate threshold.

Similar results emerged from assignment of individuals to MLLs under an infinite alleles model, using Genodive. The inferred number of MLLs (i.e., richness; Fig 4a) and effective number of MLLs (Fig. 4B) are much smaller than sample sizes for all thresholds from 14 to 28 marker differences for all patches, and they decline rapidly as thresholds are increased for grouping individuals into MLLs. All patches were inferred to belong to a single genetic lineage at greater numbers of marker differences. Patch 8 consistently was inferred to have more MLLs than other patches; for thresholds smaller than 14 the inferred number of MLLs in this patch approached the sample size, so we did not consider smaller thresholds.

**FIGURE 4.**
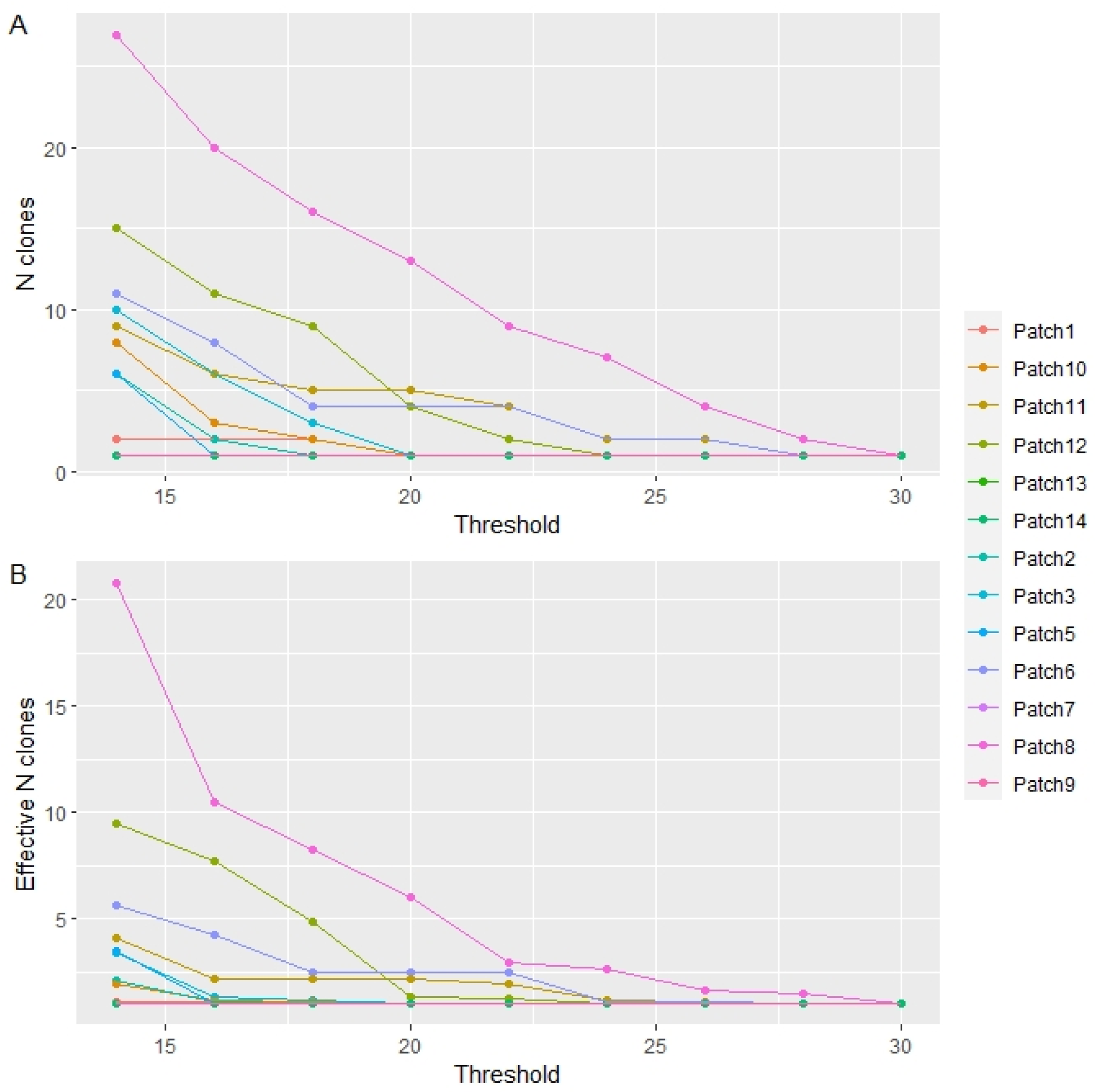
Number of MLLs per patch (A) and effective number of MLLs per patch (B), as functions of the threshold number of marker differences used to assign individuals to MLLs. Assignment was made based on the infinite alleles model.

Finally, under the assignment test (using *α* = 0.002 and 358,000 resampled individuals) no individuals were tagged as migrants.

Using the MLG dataset, the effective number of alleles at each locus is quite low (1.0-1.3) regardless of whether estimates are derived from estimates of Nei’s genetic diversity or of Shannon-Weaver allelic diversity (Table 3). Furthermore, estimates of within patch inbreeding range from 0.741 to 0.912 suggesting that genotypes are largely homozygous (Table 3). Both observations are consistent with the largely clonal structure of patches and suggest that sexual reproduction, when it occurs, largely involves close relatives or self-fertilization. In contrast, composite disequilibrium 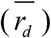 ranges between 0.015 and 0.055, and is only weakly correlated with sample size (*ρ* = 0.12; Table 3). The resampling tests suggest that the observed composite disequilibrium is not likely to reflect sampling error, but the small magnitude of the composite disequilibrium combined with the evidence for strong clonal structure suggests that variation within patches largely reflects mutations that arise independently within clonal lineages.

**TABLE 3.** Estimates of genetic diversity. *N* (number of samples, equal to the number of genotypes in the MLG dataset), allelic diversity (*H_e_*: expected heterozygosity, *n_e_*: effective number of alleles), Shannon-Weaver diversity (*H*: entropy, exp(*H*): effective number of alleles) and Inbreeding coefficients (*FIS*). Allelic diversity, Shannon-Weaver diversity and inbreeding coefficients are reported as posterior means and 95% credible intervals based on allele frequency estimates from Hickory. *r_d_* is an estimate of average composite disequilibrium in each population, and *P(r_d_)* is a test of the null hypothesis that *r_d_* = 0, based on a resampling approach.

| Patch | N | Allelic diversity |  | Shannon-Weaver diversity |  | Inbreeding | Composite disequilibrium |  |
| --- | --- | --- | --- | --- | --- | --- | --- | --- |
| | | $H_e$ | $n_e$ | $H$ | $\exp(H)$ | $F_{IS}$ | $r_d$ | $P(r_d)$ |
| 1 | 26 | 0.111<br>(0.102, 0.122) | 1.007<br>(1.000, 1.066) | 0.175<br>(0.162, 0.192) | 1.229<br>(0.162, 0.192) | 0.741<br>(0.135, 0.99) | 0.055 | 0.00 |
| 2 | 33 | 0.124<br>(0.118, 0.131) | 1.007<br>(1.000, 1.065) | 0.194<br>(0.185, 0.204) | 1.256<br>(1.244, 1.269) | 0.870<br>(0.527, 0.996) | 0.032 | 0.00 |
| 3 | 35 | 0.124<br>(0.116, 0.131) | 1.007<br>(1.000, 1.053) | 0.196<br>(0.185, 0.207) | 1.256<br>(1.241, 1.271) | 0.865<br>(0.419, 0.994) | 0.030 | 0.00 |
| 5 | 26 | 0.129<br>(0.123, 0.136) | 1.013<br>(1.000, 1.109) | 0.200<br>(0.190, 0.211) | 1.267<br>(1.253, 1.281) | 0.891<br>(0.611, 0.993) | 0.032 | 0.00 |
| 6 | 34 | 0.161<br>(0.152, 0.168) | 1.007<br>(1.000, 1.063) | 0.249<br>(0.236, 0.259) | 1.331<br>(1.313, 1.345) | 0.858<br>(0.544, 0.992) | 0.047 | 0.00 |
| 7 | 25 | 0.08<br>(0.072, 0.089) | 1.005<br>(1.000, 1.057) | 0.130<br>(0.119, 0.142) | 1.167<br>(1.151, 1.185) | 0.803<br>(0.158, 0.991) | 0.024 | 0.00 |
| 8 | 35 | 0.181<br>(0.168, 0.190) | 1.056<br>(1.002, 1.206) | 0.282<br>(0.264, 0.295) | 1.373<br>(1.347, 1.391) | 0.857<br>(0.449, 0.992) | 0.027 | 0.00 |
| 9 | 29 | 0.087<br>(0.082, 0.092) | 1.005<br>(1.000, 1.053) | 0.136<br>(0.128, 0.145) | 1.180<br>(1.169, 1.192) | 0.912<br>(0.655, 0.993) | 0.017 | 0.00 |
| 10 | 27 | 0.114<br>(0.107, 0.121) | 1.008<br>(1.000, 1.072) | 0.179<br>(0.169, 0.190) | 1.235<br>(1.222, 1.25) | 0.873<br>(0.514, 0.994) | 0.042 | 0.00 |
| 11 | 26 | 0.142<br>(0.133, 0.151) | 1.008<br>(1.000, 1.074) | 0.222<br>(0.209, 0.235) | 1.293<br>(1.275, 1.311) | 0.895<br>(0.567, 0.996) | 0.048 | 0.00 |
| 12 | 27 | 0.153<br>(0.145, 0.161) | 1.290<br>(1.063, 1.624) | 0.238<br>(0.227, 0.249) | 1.315<br>(1.300, 1.331) | 0.816<br>(0.469, 0.99) | 0.034 | 0.00 |
| 13 | 18 | 0.072<br>(0.064, 0.082) | 1.008<br>(1.000, 1.077) | 0.117<br>(0.103, 0.133) | 1.151<br>(1.134, 1.172) | 0.745<br>(0.125, 0.993) | 0.038 | 0.00 |
| 14 | 17 | 0.058<br>(0.050, 0.070) | 1.012<br>(1.000, 1.115) | 0.099<br>(0.085, 0.117) | 1.123<br>(1.107, 1.148) | 0.81<br>(0.162, 0.993) | 0.015 | 0.005 |

### Genetic structure among patches

Using Hickory, we estimated *F*_st_ to be 0.567 with a 95% credible interval of (0.519, 0.615) among patches. Using the Grunwald et al. (2016) approach to choose the optimal number of clusters, we identified 13 statistical clusters that correspond to groups of genotypes more similar to one another than to other genotypes in our sample. These clusters largely corresponded with individual patches. Principal components 1 and 2 of the PCA explained 18% of the total variation and separated individuals into groups with almost perfect correspondence between patch and statistical cluster (Fig. 5). These two axes alone separate many of the patches completely. Likewise, genetic structure is apparent from the neighbor-joining tree, which assigned all *L. chinensis* ramets into the same thirteen clusters, with almost no mixing among patches (Fig. 6). Examining the PCA assignments, we found that only in patch 8 were ramets assigned to multiple clusters; in every other patch the sampled ramets were assigned to single clusters.

**FIGURE 5.**
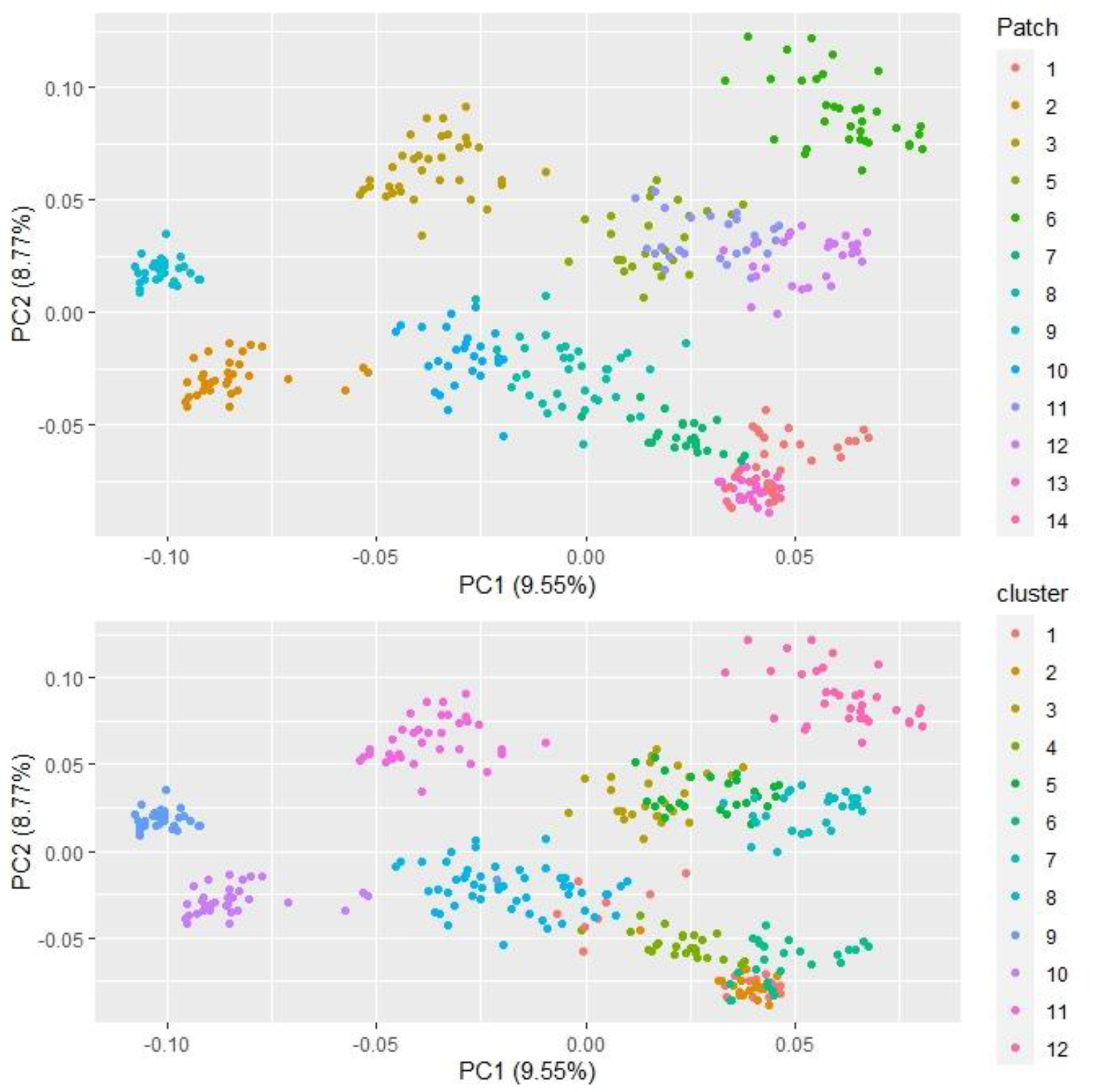
Plot of first two principal component scores. Points are labeled with (top) patch number, and (bottom) PCA cluster numbers.

**FIGURE 6.**
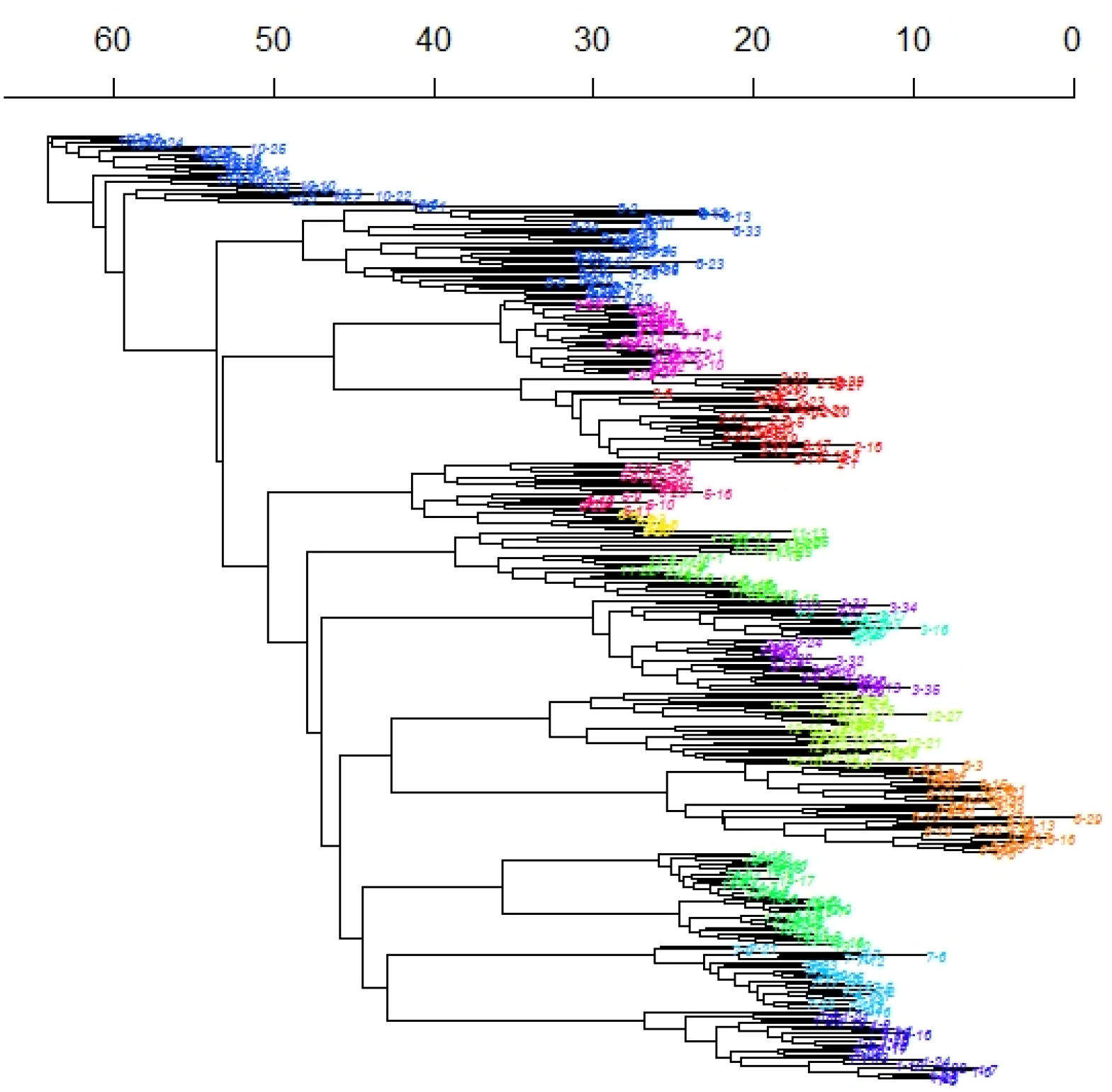
Neighbor-joining tree, with branch tips colored by cluster from the cluster analysis.

### Genetic structure within a patch

Mantel correlograms for the MLG data set found that, in some patches, over short spatial scales, there were clearly positive genetic correlations between individuals. These correlations were significant in five of the 13 patches (2, 3, 6, 11 and 14). There were no clearly negative correlations. Because only patch 8 had multiple MLLs and it had only two, analysis of MLL structure within a patch would not be informative.

### Consequences of number of markers used

Our random resampling approach to subsets of markers indicates (for the MLG data) that the relationship between number of markers used and log (sample sd of the number of genotypes identified) is strongly linear (Table 4). For 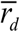, the mean of the estimates approaches its final value around 60 or so markers, but its standard deviation declines exponentially with the number of markers used (Fig. 7).

**TABLE 4.** Mean and sample variance of number of unique genotypes identified, by number of markers used.

| **Number of**<br>**markers**<br>**used** | **Mean number of**<br>**unique genotypes**<br>**identified** | **Sample variance**<br>**of number of**<br>**unique**<br>**genotypes**<br>**identified** |
| --- | --- | --- |
| 10 | 110.2 | 7044.3 |
| 30 | 303.0 | 2494.1 |
| 50 | 345.1 | 225.5 |
| 70 | 354.8 | 23.9 |
| 90 | 357.2 | 3.4 |
| 110 | 357.9 | 0.1 |
| 130 | 357.9 | 0.2 |
| 150 | 358.0 | 0.0 |

**FIGURE 7.**
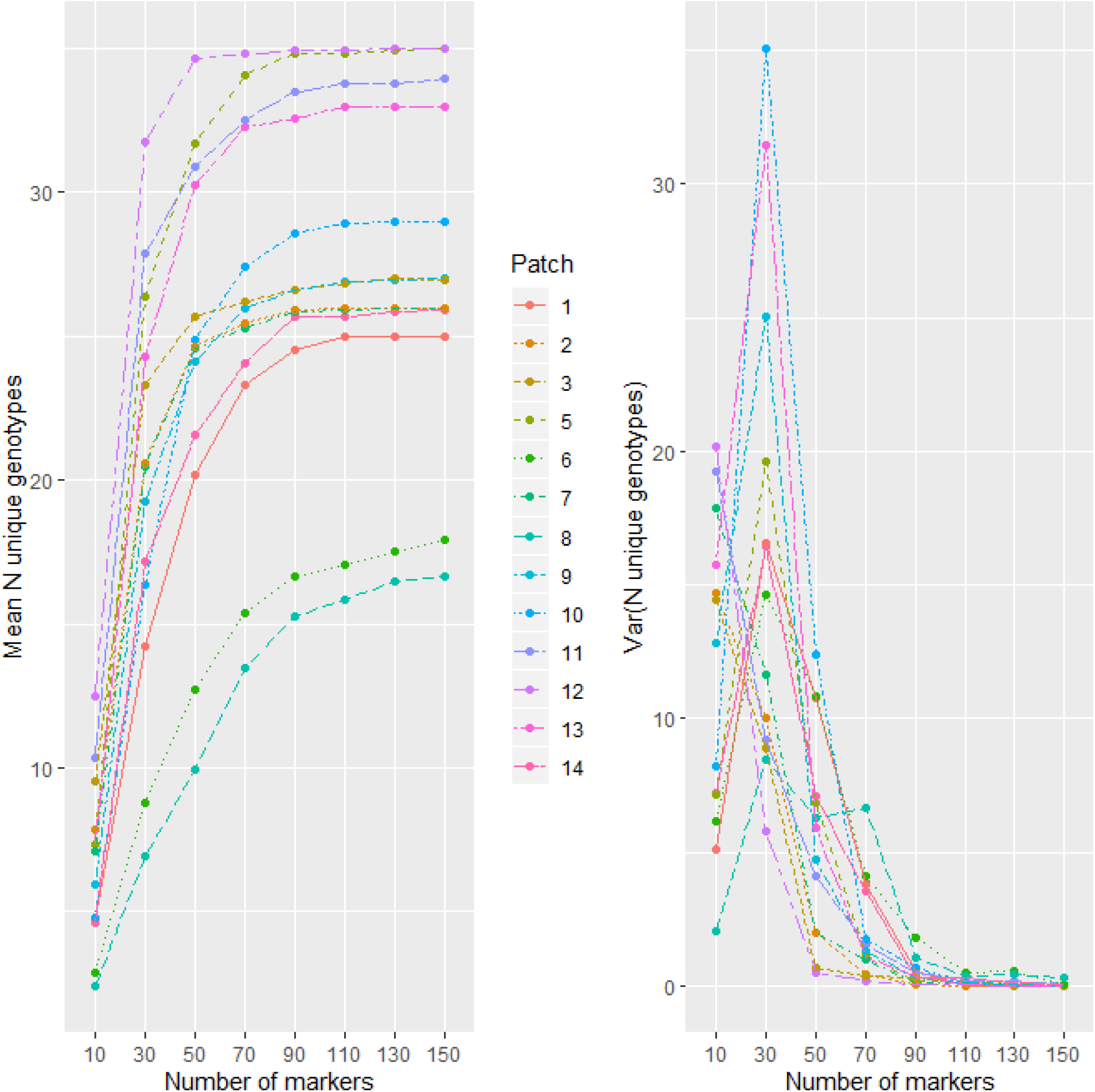
The sample means (left) and variances (right) of the number of unique genotypes in each patch, as a function of the numbers of markers used.

## DISCUSSION

Successful conservation and restoration efforts depend critically on understanding the genetic make-up of populations, which includes not only genetic diversity, but how that diversity is structured by clonal propagation within populations (Hufford and Mazer, 2003; McKay et al., 2005; Kramer and Havens, 2009). Understanding the genetic make-up of foundation plant species in particular is important for conservation since these species create and define ecological communities and contribute disproportionately to ecosystem function (Ellison et al., 2005; Hughes and Lotterhos, 2014; Lau et al., 2016; Ellison, 2019; Whitham et al., 2020). Foundation grass species often combine clonal and sexual reproduction (Franks et al., 2004; Keeler et al., 2004; Richards et al., 2004; Tumas et al., 2019). Although our initial analysis found that every ramet of the foundation grass *L. chinensis* was a unique genotype, further consideration of the reproductive biology of the species and the number and type of markers we used, suggest that we have 13 lineages that are closely associated with patches and some small amount of variation that could be due to somatic mutation as well as technical or scoring errors, which requires further study to decipher.

### Identifying unique genotypes

Every individual sampled in this study was unique (N=358; G/N = 1.0; *^1^D= ^2^D =* sample size for each patch). Several methods of identifying clusters of similar (and presumably closely related) genotypes identified 13 multilocus lineages (MLLs). Within each lineage genotypes are more closely related to each other than they are to genotypes in other lineages. Whether by identifying a minimum in pairwise Prevosti distances or by assigning individuals to lineages using an infinite alleles model, we identified approximately 13 MLLs in this study. Only one of the patches we studied contains more than one MLL identified in this way; G/N is approximately 27.5 for the entire sample, and *^1^D* and *^2^D* are close to 1 for each patch. For comparison, we used the mean value (0.62) of estimates of the Gini-Simpson index reported by Ellstrand and Roose (1987) for populations of 21 clonal species; this corresponds to an average of *^2^D =* 2.63.

The low number of MLLs found within patches is consistent with extensive clonal spread in *L. chinensis* (Yang and Li, 1994; Yang et al., 1995; Wang et al., 2005). Furthermore, very few *L. chinensis* seedlings can be found in the field (Zhu et al., 1981; Yang et al., 1995; Huang et al., 2002), even if seed production is abundant. A study in the southwestern Heilongjiang Province of China evaluated every ramet (58-79 ramets) of four plots ranging in size from 4.0-4.6 m^2^, and found only 5-15 genotypes in each patch (G/N from 0.09-0.21; Hong, 2013).

However, several studies of other clonal species found high clonal diversity. Widén et al. (1994) reported an average of 27% different unique genotypes in 45 clonal plant species based on isozyme analysis. We suggest that the difficulty may lie, at least partly, in how researchers define clones. Inferring clone membership on the basis of identical genotypes (rather than the basis of being a member of a lineage that was produced from a single sexually produced seed) can lead to confusion, because somatic mutation leads to genetic variation among ramets (Scrosati, 2002; Schoen and Schultz, 2019; Gurevitch et al., 2020). Ellstrand and Roose (1987) reported that the number of genotypes was positively correlated to the number of markers used, and was usually higher in DNA-based than in isozyme-based studies (see also Schoen and Schultz, 2019). Our results from randomly sub-sampling (Fig. 7) our data suggest that, had we used fewer markers, we might have identified many fewer unique genotypes. Studies on another perennial grass *Spartina alterniflora* reported either that nearly every ramet was unique (Richards et al., 2004; Foust et al., 2016, Robertson et al., 2017) or that the number of ramets per genet varied greatly (Hughes and Lotterhos, 2014). Moreover, Tumas et al. (2019) found that measures of genetic and clonal diversity varied dramatically across the range of the salt marsh foundation plant *Juncus roemerianus*. They found greater genetic diversity and lower clonal diversity in Gulf of Mexico populations than on the Atlantic coast. The authors suggest this could be the result of differences in plant community and disturbance regimes or reflect a relationship with population size. However, none of these studies considered the importance of closely related genets that might actually be more accurately classified as lineages. This type of analysis could inform our understanding of actual functional differences and how they might support adaptive response.

### Genetic diversity

Wang et al. (2005) examined 16 natural populations of *L. chinensis* from different geographical regions and showed that there was significant variation among populations (*F*_st_ = 0.211, *P* < 0.001). Yuan et al. (2016) investigated 18 natural populations of *L. chinensis* along a longitudinal gradient from 114 to 124°E in northeast China and concluded that genetic diversity was significantly higher in eastern (*H_exp_* = 0.628) than in western sites (*H_exp_* = 0.579) (*P* < 0.01). In this study, we found that marker richness among patches ranged from 73-134 (out of a total of 186 total markers), and our estimate of *H_exp_* ranged between 0.046 and 0.186 among patches, with an overall 0.308 (Table 2). The population-wide estimate from Hickory shows a large degree of differentiation between patches (*F*_st_ = 0.567; 95% credible interval = ((0.519, 0.615)). By contrast the pooled estimate of *H_exp_* is 0.347 (0.333, 0.362). Our among-patch differentiation thus appears to be considerably larger than in previously studied populations of *L. chinensis,* and our within-patch variation lower. The percent polymorphic loci (40-81%) among patches is on the same order of magnitude as reported for outcrossing grasses (50-60% for regional or widespread outcrossers; Hamrick and Godt, 1996; Godt and Hamrick, 1998).

Given that clonal growth is extensive and previous studies have found less than 1% survival rate of seedlings in *L. chinensis* natural populations (Benson et al., 2004; Dalgleish and Hartnett, 2006), it is likely that the high level of genetic variation could be due to somatic mutation, or occasional cross pollination, and long life span. Somatic mutation would create unique loci and contribute to the high diversity observed within patches of *L. chinensis* (Antolin and Strobeck, 1985). Lynch (1984) and Gill et al. (1995) both reported that high rates of somatic mutation could compensate for the absence of recombination from sexual reproduction in primarily asexual species, and may allow asexual species to maintain abundant genetic variation and adapt to changing environmental conditions. However, the mutation rate in plants in general is poorly understood (Schoen and Schultz, 2019). This is also true among clonal plant species, despite the fact that several authors have argued the importance of somatic mutation in clonal plants (Eckert, 2002; Fischer and van Kleunen, 2002). Somatic mutation has generally been considered disadvantageous with the expectation that there should be selection for low somatic mutation rates. On the contrary, in a recent review of the impacts of somatic mutations in evolution Schoen and Schultz (2019) argue that in plants, “somatic mutation might be relatively innocuous or even beneficial, as the indeterminate and partially independent modular design of most plants may allow cell lines to be lost without sacrificing overall plant function and can amplify the occurrence of beneficial mutations.” They also present evidence that the contributions of somatic mutation to population genetic variability varies in different species. For instance, high genotypic diversity in Australian populations of *Grevillea rhizomatosa* (Proteaceae) was attributed to somatic mutations and 25% of connected ramets had differences based on ISSR markers (Gross et al., 2012). Similarly, in New Zealand populations of *Hieracium pilosella*, high diversity of ISSR genotypes could have resulted from somatic mutation (Houliston and Chapman, 2004).

In our study, somatic mutation might be indicated by markers that are only found in a few individuals or if pairwise comparisons indicated that many ramets were differentiated by only a few markers. However, we found that only eight markers (2%) were found in only one or two ramets. Moreover, more than 99% of ramets differed by more than 10 markers. Then again, these expectations might not be reasonable depending on the age of the patches and the accumulation of multiple mutations within MLLs. Further work is required to identify the importance of somatic mutation in these populations.

Although the capacity for sexual reproduction of *L. chinensis* is reportedly rather weak, frequent somatic mutations and some level of sexual reproduction could generate considerable variation. These loci would be accumulated through extensive clonal propagation and thus, each patch would exhibit a high level of genetic diversity. We evaluated the number of markers that would be effective at identifying the minimum number of MLL in this study to be 60 or so markers instead of the 186 that we used here. We suggest further study will be required to evaluate how many markers to use, and how to interpret the resulting data, given the possible maintenance of somatic mutations in highly clonal plants. The contribution of somatic mutation could be an important consideration for markers like microsatellites that already have high expected mutation rates.

### Genetic structure among patches

Clear and strong genetic structure among the thirteen patches were observed both in PCA plots and neighbor-joining trees providing evidence for genetic isolation among these patches, and a lack of gene flow among patches. Gene flow can occur at several life stages in plant species: as pollen, fertilized seeds or rhizome movement via water or animal vectors or disturbance. Previous studies report low pollen viability, short pollen longevity and short pistil receptivity of *L. chinensis* lead to low seed production (Huang et al., 2004). The *L. chinensis* seeds do not have wings or plumes for dispersal by wind. Thus, lack of dispersal ability could result in a decreased possibility of sexually produced genetic exchange. In addition, isolation could result from the history of colonization of a patch by specific genotypes with subsequent accumulation of mutations.

If somatic mutation is an important source of variation, the differences among patches could reflect an age-based structure as mutations accumulate over time. However, a study of the full genome of a single 234-year-old European oak (*Quercus robur*), found 17 point mutations while another study found almost three times as many point mutations in an even younger tree (Schmid-Siegert et al., 2017; Plomion et al., 2018). Up to seven out of 19 mutations were also transmitted to offspring indicating inheritance of these differences not only through vegetative reproduction but through sexual (Plomion et al., 2018). In a study of aspen, Ally et al. (2010) found that the number of somatic mutations at 14 microsatellite loci was a good proxy for individual plant age, while plant (clone) size alone was not a reliable indicator.

Finally, there may be spatial variation of selection intensity induced by environmental heterogeneity (Linhart and Grant, 1996). Owing to microhabitat heterogeneity, alleles within plant populations can have a strong spatial structure (Hamrick and Holden, 1979). If this process is partly responsible for the variation among patches, it would be operating because of linkage between ISSR loci and loci under selection. Thus inferences about it would require much more information about the *Leymus* genome as well as about spatially heterogenous selection. Identifying the contribution of limited migration, somatic mutation, age and microhabitat heterogeneity to the genetic structure among patches of *L. chinensis* will require more detailed studies of these populations and environmental factors.

### Genetic structure within a patch

We detected significant and positive correlations between geographic and genetic distances among individuals within either the three smallest patches (11, 13, 14) or the four largest patches (2, 3, 6, 8) with Mantel tests, indicating that geographic distance is a factor shaping the genetic structure of these patches. Yang and Zhang (2006) reported that with increasing patch area, the number of total ramets increased linearly in *L. chinensis* patches. The clonal architecture of *L. chinensis* tended to be spatially clustered in patches 13 and 14. In this case, neighboring ramets are expected to be closely related, such that positive autocorrelation may exist within a short distance. Clonal growth form can greatly influence the distribution of genetic variation within clonal patch.

For the largest patches (2, 3, 6 and 8), low ramet densities and larger spatial distances between ramets, may have further restricted gene flow among individuals. The observed negative correlations over the longest distance in patches 6 and 8 indicated genetic differentiation within a patch may not occur in patches smaller than 8 meters in diameter. In summary, patch size plays an important role in shaping the genetic structure among individuals within patches.

### Implications for conservation

Genetic diversity is the material basis for the existence and development of populations, and protecting genetic diversity of populations is an important task. Several authors acknowledge that while foundation species are common and abundant, and therefore often not considered for conservation purposes, protecting these species is critical for maintaining ecosystem function (Ellison, 2019; Qiao et al., 2021). *Leymus chinensis* is a candidate foundation species in the Inner Mongolia steppe region, where overgrazing has led to severe degradation (Bai et al., 2004; Tong et al., 2004). In addition, these grasslands face extreme climate events and chronic environmental challenges that are partly buffered by this species (Bai et al., 2010; Meng et al., 2019). Restoration of these degraded populations could depend on management of patchy populations. In order to do so, it is necessary to understand how genetic diversity is structured within and among patches before formulating a conservation or restoration strategy, bearing in mind that a recent study reported significant variation among 10 *L. chinensis* genotypes in response to drought, high temperature, and both drought and high temperature (Yang et al., 2019). Our results indicated larger patches have higher levels of genetic diversity, but patches themselves are frequently derived from a single lineage. These findings provide important reference information for management agencies. Although we do not know how functionally different these MLLs are, we suggest that the conservation management of these patchy populations should keep to the following principles: if conditions permit, it is better to protect as many patches as possible; if resources are limited, local managers should give priority to protecting several large patches because of their high genetic diversity, and increased potential for functional diversity. Maximizing the retention of genetic diversity will be important for the long-term survival of patchy populations.

## ACKNOWLEDGMENTS

The authors thank Qian Xiong for assistance with the experiment. The study was financially supported by National Science Foundation of China (Grant No. 32071860 and 31570332), Natural Science Foundation of Science and Technology Department of Liaoning Province (Grant No. 2019-MS-155), Serving Local Project of Education Department of Liaoning Province (Grant No. LFW202001), and Federal Ministry of Education and Research (BMBF; MOPGA Project ID 306055 to CLR). The authors gratefully acknowledge financial support from China Scholarship Council.

## AUTHOR CONTRIBUTIONS

The project was designed and organized by Chan Zhou. Jian Guo and Zhuo Zhang conducted and managed data collection. Gordon A. Fox coordinated statistical analysis. Zhuo Zhang provided critical molecular expertise throughout the project. Kent E. Holsinger and Christina L. Richards provided expertise in population genetics. All authors contributed to the writing of the manuscript.

## DATA AVAILABILITY

Data and code for data analysis will be archived with Datadryad (http://datadryad.org) upon acceptance of the paper.

